# Ancestry-specific gene expression in peripheral monocytes mediates risk of neurodegenerative disease

**DOI:** 10.1101/2024.11.20.624489

**Authors:** Aaron Z Wagen, Regina H Reynolds, Jia Nee Foo, Aine Fairbrother-Browne, Emil K Gustavsson, Sarah Galgiano-Turin, Nicholas W Wood, Cornelis Blauwendraat, Sonia Gandhi, Mina Ryten

## Abstract

It is hypothesised that peripheral immune states responding to regional environmental triggers contribute to central neurodegeneration. Region-specific genetic selection pressures require this hypothesis to be assessed in an ancestry specific manner. Here we utilise genome-wide association studies and expression quantitative trait loci from African, East Asian and European ancestries to show that genes causing neurodegeneration are preferentially expressed in innate rather than adaptive immune cells, and that expression of these genes mediates the risk of neurodegenerative disease in monocytes in an ancestry-specific manner.

## Main

The role of ancestry-specific risk is increasingly recognised in neurodegenerative diseases (NDDs). Rectifying the historical focus on European populations, recent efforts have explored the clinical and genetic diversity of conditions such as Alzheimer’s (AD) and Parkinson’s diseases (PD), and Frontotemporal dementia (FTD)^1–4^. This includes appreciation of the differences in the prevalence and pathology of NDDs across populations^5,6^, as well as differences in the genetic architecture of both monogenic and complex forms of disease^3,4,7,8^. Central to this has been the discovery of novel disease loci in ancestry-specific genome-wide association studies (GWAS) of NDDs^7–11^, and of variability of gene expression in post-mortem brains^6,12^. Despite these studies being limited by relatively small case numbers, they have identified novel risk loci with unexpectedly large effect sizes, emphasising the significant effect ancestry can have in neurodegeneration.

Simultaneously, the immune system is increasingly implicated in NDDs, including AD, PD and FTD. Within the CNS, the role of microglia, the parenchymal myeloid cells of the CNS, as well as immune related macro-glial states are gaining prominence^13–15^. Furthermore, both positron emission tomography (PET) based imaging approaches, as well as cerebrospinal fluid (CSF) biochemical analyses have demonstrated increases in inflammatory biomarkers in NDDs, with levels correlated to clinical outcomes. This central immune response is mirrored peripherally, and there is suggestive evidence that the peripheral response is fundamentally involved in disease pathogenesis^16,17^. Clinically, NDDs can commence with a peripheral prodrome, for example, anosmia and constipation in PD. Epidemiological data shows that NDDs have followed viral pandemics throughout history^18^, and that recent viral infection increases the risk of NDD^19,20^. Interestingly, this appears to occur without evidence of replicating viral particles, raising the possibility of a virally-triggered peripheral immune state that affects CNS function. Indeed, the concept of brain-immune crosstalk has been advanced with the potential to explain how environmental factors might contribute to neurodegenerative risk^21^, and underpinning recent therapeutic strategies to treat these diseases^22–24^.

In this work, we hypothesise that genes causally linked to AD, PD and FTD are themselves active in the peripheral immune response, and that this activity contributes to the risk of disease in an ancestry-specific manner. There is already some evidence to support this idea in PD, where studies in mice have shown that alpha-synuclein (*SNCA*), the major constituent of Lewy bodies and a cause of PD, has a protective role against viral infection centrally^25^, and can mediate antigen presentation and the inflammatory response in the peritoneum^26^.

Similarly, common pathogenic variants in leucine-rich repeat kinase-2 (*LRRK2*) modulate inflammation in response to CNS and systemic infections^27,28^, as does a pathogenic *GBA1* variant with respect to viral encephalitis^29^. This association is also seen in AD and FTD, where, respectively, *TREM2*^30,31^ and *C9orf72*^32^ have peripheral immunomodulatory effects. Genetic analyses have linked the heritability of NDDs with peripheral immune cell types^33,34^. However, the risk of NDDs caused by the expression of genes in peripheral immune cells across diverse ancestries has not been investigated.

To study this, we systematically explored the role of 39 NDD genes causally implicated in AD, PD or FTD across peripheral immune cells (Supplementary Table 1, Methods). Given that both ancestry and activation state are well recognised to affect gene expression in peripheral immune cells, we used a multi-ancestry study of 22 different cell-types to study gene expression^35^. This dataset was derived from 222 African (n = 82), East Asian (n = 60) and European (n = 80) donors, either at baseline or after activation with SARS-CoV-2 or Influenza A viruses.

We found that 32 of the 39 NDD-causing genes (82%) were expressed across peripheral immune cells (Figure 1a). Using phenotypic QTL data, we noted that 29 of these 32 genes (91%) were also causally associated with a white blood cell count metric in peripheral blood. Indeed, 20-83% of the NDD-causing genes, including *APOE, SNCA* and *MAPT*, were loci that influenced macrophage, monocyte, lymphocyte or total white cell count in at least one ancestry (Figure 1b, Supplementary Figure 1).

**Fig. 1:**
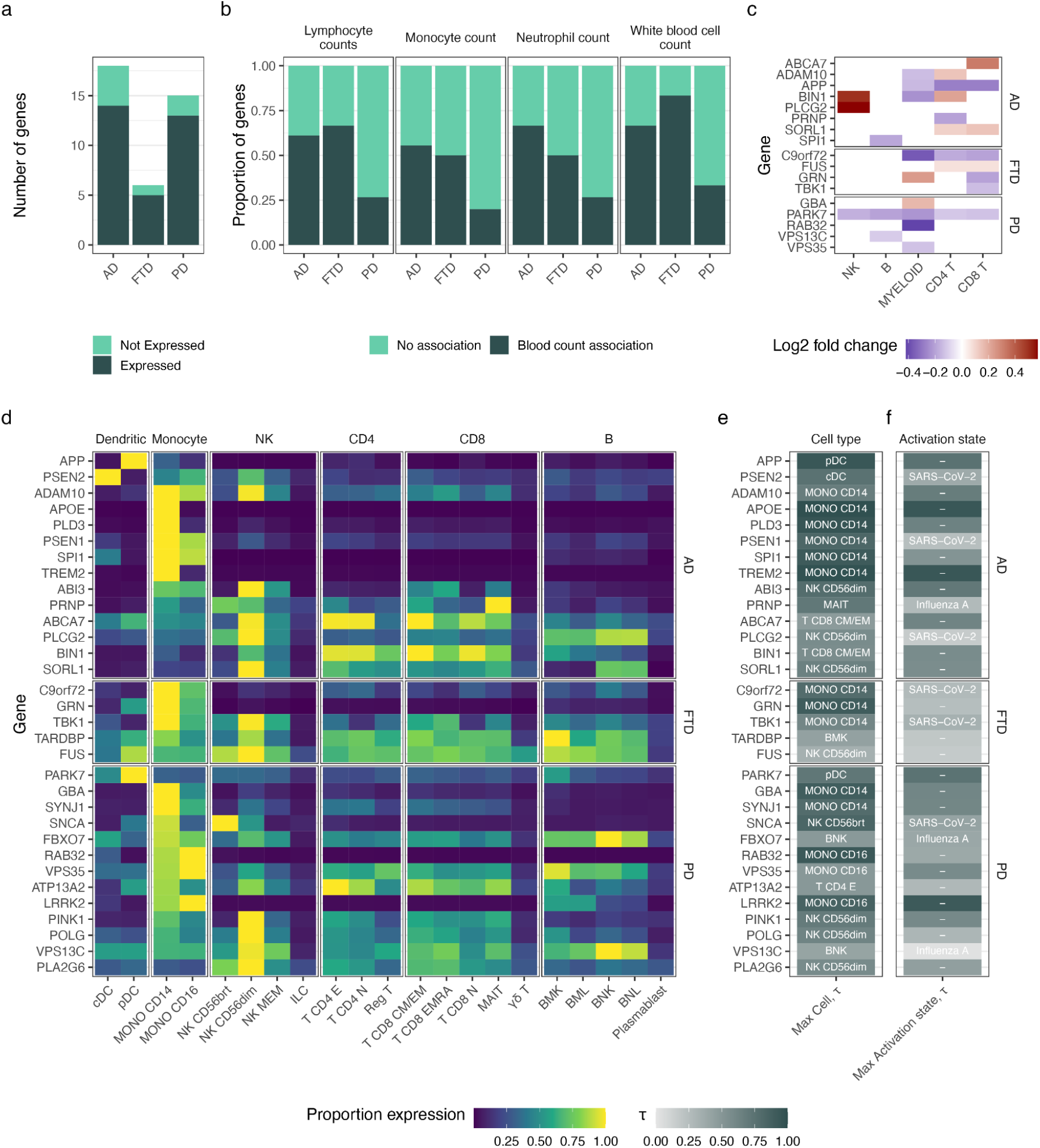
Genes causing neurodegeneration are expressed in peripheral immune cells in a cell-type, cell-state and ancestry specific manner. **a**, Number of genes causing neurodegeneration expressed in multi-ancestry study of peripheral blood cells (Aquino et al^35^). **b**, Proportion of genes expressed in peripheral blood cells (14 AD, 13 PD and 5 FTD) that have an association with lymphocyte, monocyte, neutrophil and total white blood cell count (left-to-right, from Chen et al^53^). **c**, Results of differential gene expression analysis, showing genes significantly differentially expressed in blood cell lineages in European relative to African ancestry. Empty tile denotes no significant differential expression. **d** Heatmap showing proportion of gene expression of genes causing neurodegeneration across cell types and lineages in multi-ancestry study. The cell-state with maximal gene expression is shown, with expression represented as proportion of maximal expression per gene over all cell types. **e**, Column showing cell-type specificity of genes, with labels showing the cell-type with greatest gene expression, and shaded by τ-statistic (darker suggests greater specificity). **f**, Column showing cell-state specificity, where, for the cell-type that showed maximal expression for each gene, the label shows the exposure that caused the maximal expression (or ‘-’ where maximal expression was in unstimulated cells). Shading shows the τ-statistic of the labelled activation state compared to the other activations.

Next, we explored the expression patterns of NDD-causing genes by cell type, activation state, and ancestry. Over 80% of these genes showed maximal expression in innate immune cell types, most commonly in monocytes (16 out of 34 cell types) and NK cells (8 out of 34 cell types, Figure 1d). Genes with maximal expression in myeloid cell types (including *APOE, PSEN1, TREM2, LRRK2* and *C9orf72* in monocytes; and *APP* and *PSEN2* in dendritic cells) showed greatest cell-type specificity (mean τ-statistic = 0.82, Figure 1e). In contrast, genes with maximal expression in NK cells were more broadly expressed (mean τ-statistic 0.63, p value for two sample T-test = 0.01). Three NDD genes (*TREM2, APOE* and *LRRK2*) showed high specificity (τ-statistic > 0.9) to unstimulated activation states as compared to those treated with influenza or SARS-CoV-2 (Figure 1f). Notably, these three genes all had maximal expression in monocytes. Myeloid cells also showed the greatest number of genes differentially expressed across ancestries (European relative to African ancestry) with 10 genes identified, including *APP, BIN1, GBA1, RAB32, C9orf72* and *GRN* (Figure 1c). Taken together, these results show that genes central to NDDs are preferentially expressed in peripheral immune cells of myeloid lineage, and that these cells contain the greatest proportion of cell type- and ancestry-specific NDD gene expression.

We then explored whether the expression of these genes in the periphery might be causally linked with disease. To do this, we applied a well-established genetic approach, colocalisation analysis, to explore the possibility that NDD gene expression in the peripheral immune system might be a driver of disease risk^36^. This analysis uses a Bayesian approach to ascertain the posterior probabilities that association signals of two traits at a genetic locus share a causal variant (posterior probability of hypothesis 4, PPH4), or whether these association signals are distinct (posterior probability of hypothesis 3, PPH3, Methods).

Crucially, these analyses were conducted in an ancestry-specific manner, as we reasoned that the peripheral immune system would be the first to encounter geographically restricted microbes and pollutants, leading to variable environmental selection pressures^37^. This required use of very recently produced ancestry-specific GWAS, as well as corresponding ancestry-aware functional genomic annotations, namely expression quantitative trait loci (eQTL). We utilised the eQTL available from the same multi-ancestry study used to explore gene expression above, due to the availability of data across multiple populations, cell types and activation states^35^.

We first explored ancestry-specific AD GWASs with multi-ancestry eQTL data. When using a European AD GWAS^38^, we identified a colocalising signal in *BIN1* in monocytes (PPH4 = 0.93, Supplementary Table 2, Supplementary Fig. 2). There was a positive correlation between effect sizes of the GWAS and eQTL single nucleotide polymorphisms (SNPs) at this locus, suggesting that increased *BIN1* expression in peripheral monocytes correlated with increased risk of AD. While the role of *BIN1* in modulating AD is known within the CNS^39^, this result suggests a causative role for the gene in AD the periphery. Noting that *BIN1* is not a significant locus in the East Asian AD GWAS^11^, we found no colocalising signal at BIN1 when this GWAS was tested against the multi-ancestry eQTL (Supplementary Table 3).

**Fig 2.**
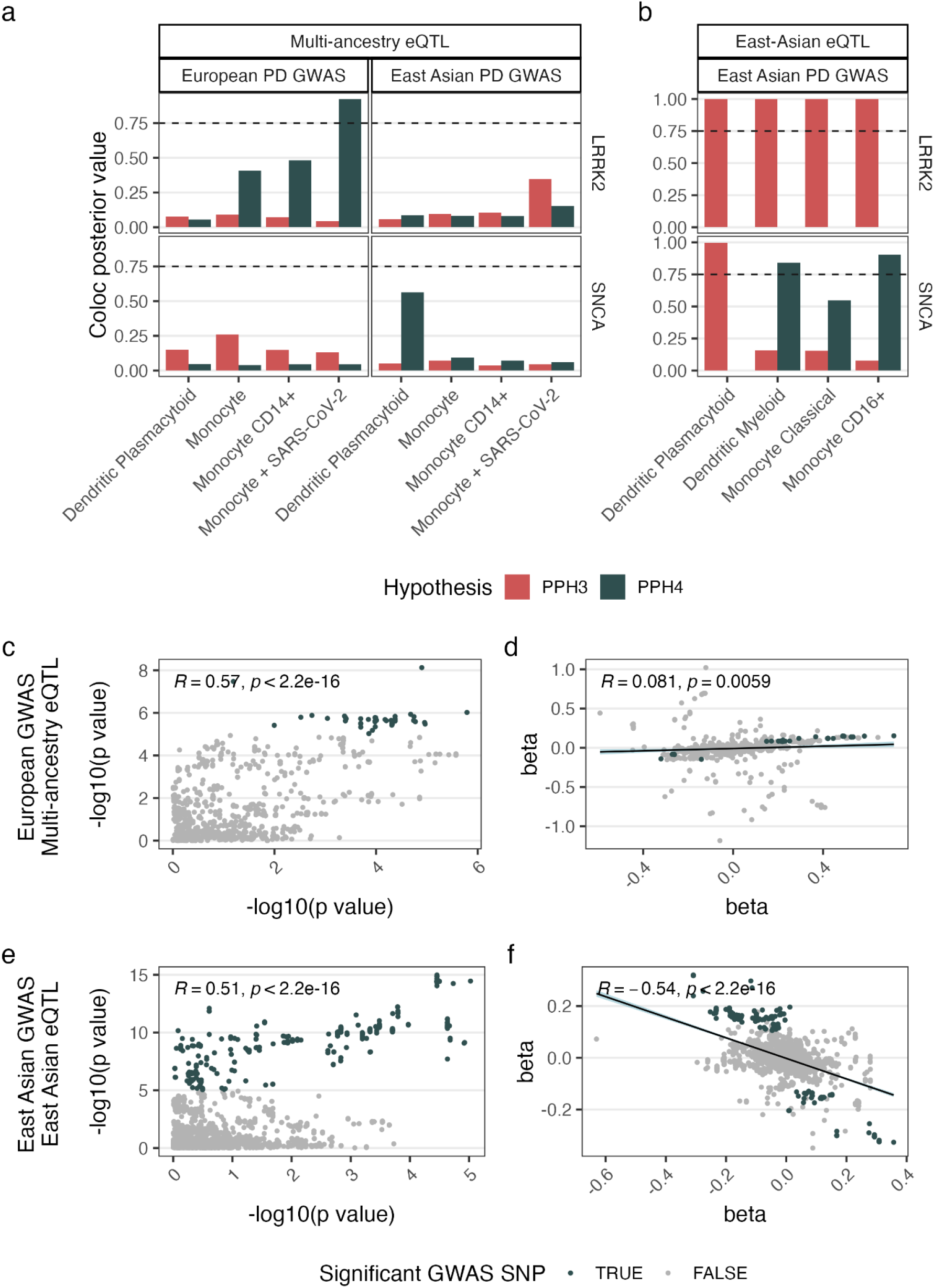
Expression of PD genes in peripheral immune cells mediate risk of PD in an ancestry dependent manner. **a**, Colocalisation of *LRRK2* (top) and *SNCA* (bottom) between multi-ancestry eQTL and PD GWAS from European (left) and East Asian (right) ancestry. PPH3 measures discrete risk, PPH4 measures colocalised risk (significant colocalisation = posterior hypothesis > 0.75, represented by the dashed line). **b**, Colocalisation of *SNCA* using eQTL and GWAS from East Asian ancestry. **c-f**, Locus analysis of significant colocalising signals between PD GWASs and eQTLs. Results from LRRK2 locus using European PD GWAS and multi-ancestry eQTL in SARS-CoV-2 treated monocytes, showing -log10(p values) (**c**) and betas (**d**). Results from *SNCA* locus using East Asian PD GWAS and East Asian eQTL in CD16+ monocytes, showing correlation of p-values (**e**) and beta values (**f**). Text shows Pearson’s correlation coefficient and p value, with beta plots also showing the regression line of best fit shaded with 95% confidence intervals.

Focusing on PD, with a GWAS derived from individuals of European ancestry^40^, we identified a colocalising signal within the PD-associated gene *LRRK2*, mediating the risk of PD and gene expression in SARS-CoV-2-treated CD14 monocytes (Figure 2a, Supplementary Table 4, Supplementary Figure 3). To check whether this result was ancestry specific, we undertook the same analysis with an East Asian PD GWAS^9^, finding that the result was underpowered at this locus (PPH3 + PPH4 < 0.75, Figure 2a, Supplementary Table 5, Supplementary Figure 4, Methods). Given this, we checked the East Asian GWAS against an East-Asian-specific eQTL dataset^41^, finding the association signals underlying *LRRK2* expression in monocytes were distinct from those underlying PD risk in all monocyte cell types tested (PPH3 >0.99, Supplementary Table 6, Supplementary Figure 5).

Focusing on the *SNCA* locus, arguably the most important gene in PD, there was insufficient power when the multi-ancestry eQTL was tested against both the European and East Asian PD GWASs (PPH4 + PPH3 < 0.75, Figure 2a, Supplementary Table 4, 5). However, noting that the PPH4/PPH3 ratio was high at 3.5, we checked this locus using an East-Asian-specific PD eQTL and GWAS. We found that in this East-Asian-specific analysis that there was a significant colocalisation in *SNCA* expression in CD16+ monocytes (Figure 2b, Supplementary Table 6, Supplementary Figure 5). Using the multi-ancestry eQTL with an African PD GWAS, there was a suggestive but underpowered colocalising signal at BIN1 in CD14+ monocytes (PPH4=0.71, PPH3+PPH4=0.73, PPH4/PPH3 ratio=39.9). We could not validate this result in an African-specific eQTL dataset given the lack of power in the available datasets (Supplementary Table 8).

As expected, at the *LRRK2* locus, there was a positive correlation between effect sizes in European populations, implying that increased *LRRK2* expression associates with increased PD risk (Figure 2b). In contrast,at the *SNCA* locus in East Asian populations, the correlation between effect sizes was negative, suggesting that decreased *SNCA* expression in CD16+ monocytes was associated with increased PD risk. While this result appears to be counter-intuitive given that increased *SNCA* in the CNS is associated with PD, this finding is in keeping with multiple existing genomic and proteomic studies, which show that PD is associated with a decreased *SNCA* RNA and protein in blood and CSF^33,42–44^.

In summary, we show that genes causally implicated in NDDs mediate the risk of these diseases partially through effects in the peripheral immune system. Crucially, the expression of these genes as well as the risk mediated by them were ancestry dependent. This suggests not only that neurodegeneration has a peripheral component, but that across different ancestries regional selection pressures may have differentially modulated disease mechanisms. The results also suggest that genes that have a detrimental effect within the CNS may be protective peripherally^25,26^. In addition to emphasising the importance of ancestry-specific functional genetic annotations, this work defines innate immune cells, and specifically monocytes, as potential targets for treatments for neurodegeneration.

## Methods

### Neurodegenerative gene list curation

Genes causally implicated in AD were identified as described by Neuner et al^45^, selecting the 18 genes with increased confidence of causation. Fourteen genes causally implicated in PD with high or very high confidence were selected as reported by Blauwendraat et al^46^, with the addition of *RAB32* whose role in PD has recently emerged^47,48^. Causative FTD genes were selected from Antionioni et al^49^, including the 6 genes that contribute ≥1% of disease frequency.

### eQTL preprocessing

We systematically searched PubMed for eQTL studies of peripheral blood from the last 10 years with the following features: i) included more than 150 participants; ii) specifically stated ancestry; iii) divided cells into clearly defined subtypes by single cell sequencing or cytometric approaches; and iv) included an activation state generated either with an immunogenic trigger or through the inclusion of participants with immunological disease. The Aquino et al dataset^35^ was the only dataset identified that included participants from African, Asian and European ancestry. Noting that it analysed 22 different cell types, and included activation state data with samples stimulated with either SARS-CoV-2 virus or influenza A virus, it was selected as the baseline dataset for this study. For colocalisation analyses, the Aquino et al dataset was supplemented by use of Ota et al^41^, as a large dataset specific to East Asian ancestry. The only African-specific ancestry eQTLs that were identified included two datasets from African and African admixed populations, one exploring monocytes^50^, and the other macrophages^51^.

Where relevant, summary statistics were processed by lifting over genomic locations from GRCh37 to GRCh38. SNP genomic coordinates were mapped to Reference SNP cluster IDs (rsIDs) using the SNPlocs.Hsapiens.dbSNP144.GRCh38 package, and ancestry-specific minor allele frequencies were imported from the MafDb.1Kgenomes.phase3.GRCh38 package^52^.

### Blood count phenotype-QTL processing

Significant associations between genetic variation and blood count metrics were derived from a multi-ancestry phenotype QTL^53^. Significant traits between blood cell metrics of interest (neutrophil, lymphocyte, monocyte and total white blood cell count) and NDD genes were exported using Open Targets (https://genetics.opentargets.org/, RRID:SCR_014622)^54^.

### Gene expression analysis

The gene expression and differential gene expression datasets used in this study were generated by Aquino et al^35^. To facilitate comparison between cell types, the raw transcript per million (TPM) for each gene was normalised by the maximum TPM per gene across cell types and activation states, giving the metric of proportion of expression per gene. Noting that cells in Aquino et al were treated with both influenza A and SARS-CoV-2 viruses, this was also normalised using the above approach. Of each cell type that showed maximal expression of a gene, the maximal cell state was defined as the viral exposure that resulted in the maximal expression of the gene for that celltype.

To measure the cell-type specificity of gene expression, the τ-statistic was calculated using the method defined by Yanai et al^55^:

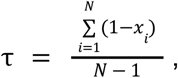

where N is the number of tissues and x_i_ is the expression profile component normalised by the maximal component value. For each cell type that contained the highest expression per gene, a further τ-statistic was calculated to measure cell-state specificity of response to viral stimulation.

Differential gene expression analysis results were utilised from Aquino et al. They compared European relative to African populations using a linear regression model, adjusted for individual donor age, cellular mortality and cellular composition as described^35^.

### GWAS preprocessing

GWAS summary statistics from 3 ancestry-specific studies of PD were used: European^40^, African^10^, and East Asian^9^ ancestry. Two ancestry-specific GWAS studies were used for AD: European^38^ and Asian^11^. Where relevant, summary statistics were processed by lifting over genomic locations from GRCh37 to GRCh38. SNP genomic coordinates were mapped to rsIDs using the SNPlocs.Hsapiens.dbSNP144.GRCh38 package (https://bioconductor.org/packages/release/data/annotation/html/SNPlocs.Hsapiens.dbSNP144.GRCh38.html)^56^, and ancestry-specific minor allele frequencies were imported from the MafDb.1Kgenomes.phase3.GRCh38 package (https://www.bioconductor.org/packages/release/data/annotation/html/MafDb.1Kgenomes.phase3.GRCh38.html)^52^.

### Colocalisation analysis

To evaluate the probability that GWAS loci and eQTLs share a single causal variant, a colocalisation analysis was performed using coloc (version 5.1.0.1, https://cran.r-project.org/package=coloc) and colochelpR (version 0.99.1, http://dx.doi.org/10.5281/zenodo.5011869)^36,57^. GWAS loci within 1 Mb significant GWAS SNPs were explored. The prior probability that any random SNP in the region is associated with the GWAS (p_1_) or eQTL (p_2_) was set to the default 10^−4^, whereas the prior probability that any random SNP in the region is associated with both traits (p_12_) was set to 10^−5^. Using a Bayesian approach, posterior probabilities were calculated for four different hypotheses that together sum to 1. Hypothesis 1 and 2 measure associations of the GWAS locus or the eQTL locus, where there is no sufficient power in the two studies to compare the two (as defined by PPH4 + PPH3 < 0.75). Hypothesis 3 (PPH3) measures the probability that the traits have distinct causal variants at a locus, while hypothesis 4 (PPH4) measures the probability that a locus is colocalised as a result of a single causal variant.

### Statistical analysis

All statistical analysis was undertaken in R version 4.2.0 (RRID:SCR_001905, https://www.r-project.org/).

## Code availability

The complete code utilised in this work can be accessed at https://github.com/aaronwagen/peripheral_immune_neurodegeneration, including knitted markdowns with integrated analysis, results, and packages utilised.

## Data availability

Summary statistics for Foo et al, Rizig et al and Shigemizu et al were supplied by the authors. Otherwise, all datasets utilised are from public sources (Supplementary Table 9).

## Acknowledgements

This research was funded in part by Aligning Science Across Parkinson’s [ASAP-000509 and ASAP-000463] through the Michael J. Fox Foundation for Parkinson’s Research (MJFF). This work utilized the computational resources of the NIH HPC Biowulf cluster (http://hpc.nih.gov). This work was supported in part by the Intramural Research Program of the National Institutes of Health including: the Center for Alzheimer’s and Related Dementias, within the Intramural Research Program of the National Institute on Aging and the National Institute of Neurological Disorders and Stroke.

AZW was supported through the award of a Clinical Research Fellowship funded by the Wolfson Foundation and Eisai Ltd. SG was supported by Wellcome (100172/Z/12/2) and is currently an MRC Senior Clinical Fellow (MR/T008199/1). MR was supported by the UK Medical Research Council (MRC) through her award of Tenure-track Clinician Scientist Fellowship (MR/N008324/1).

