## Supplementary materials for "Ancestry-specific gene expression in peripheral monocytes mediates risk of neurodegenerative disease"

### Supplementary Tables

#### Supplementary Table 1

| Disease | Gene |
|---|---|
| AD | ABCA7 |
| AD | ABI3 |
| AD | ADAM10 |
| AD | APOE |
| AD | APP |
| AD | BIN1 |
| AD | CD33 |
| AD | CR1 |
| AD | NOTCH3 |
| AD | PLCG2 |
| AD | PLD3 |
| AD | PRNP |
| AD | PSEN1 |
| AD | PSEN2 |
| AD | SORL1 |
| AD | SPI1 |
| AD | TREM2 |
| AD | UNC5C |
| PD | ATP13A2 |
| PD | DNAJC6 |
| PD | FBXO7 |
| PD | GBA |
| PD | LRRK2 |
| PD | PARK7 |
| PD | PINK1 |
| PD | PLA2G6 |
| PD | POLG |
| PD | PRKN |
| PD | RAB32 |
| PD | SNCA |
| PD | SYNJ1 |
| PD | VPS13C |
| PD | VPS35 |
| FTD | C9orf72 |
| FTD | FUS |
| FTD | GRN |
| FTD | MAPT |
| FTD | TARDBP |
| FTD | TBK1 |

**Supplementary Table 1: Table of genes causally implicated in Neurodegenerative diseases.** Alzheimer's disease (AD) genes were derived from Neuner et al[1], Parkinson's disease (PD) genes from Blauwendraat et al[2], and Frontotemporal dementia (FTD) genes from Antonioni et al[3].

#### Supplementary Table 9

| Data Resource | Link | Notes |
|---|---|---|
| Antonioni et al[3] | https://doi.org/10.3390/ijms241411732 | Data taken from tables and supplementary tables. |
| Aquino et al[4] | https://doi.org/10.1038/s41586-023-06422-9 | Data in Figure 1 taken from tables and supplementary tables.<br>Data for Figure 2 downloaded from https://dataset.owey.io/doi/10.48802/owey.e4qn-9190 on 6th April 2024. |
| Blauwendraat et al[2] | https://doi.org/10.1016/S1474-4422(19)30287-X | Data taken from tables and supplementary tables. |
| Chen et al[5] | https://doi.org/10.1016/j.cell.2020.06.045 | Downloaded using the opentargets portal on 9th September 2024. |
| Foo et al[6] | https://doi.org/10.1001/jamaneurol.2020.0428 | Summary statistics provided by authors. |
| Jansen et al[7] | https://doi.org/10.1038/s41588-018-0311-9 | Summary statistics downloaded from https://cncr.nl/research/summary_statistics/ on 12th May 2024. |
| Nalls et al[8] | https://doi.org/10.1016/S1474-4422(19)30320-5 | Downloaded from https://pdgenetics.org/resources, excluding 23andMe cases. |
| Nedelec et al[9] | https://doi.org/10.1016/j.cell.2016.09.025 | QTD000379.all.tsv.gz, QTD000384.all.tsv.gz and QTD000389.all.tsv.gz downloaded from https://www.ebi.ac.uk/eqtl/ on 13th March 2024. |
| Neuner et al[1] | https://doi.org/10.1016/j.nbd.2020.104976 | Data taken from tables and supplementary tables. |
| Ota et al[10] | https://doi.org/10.1016/j.cell.2021.03.056 | Downloaded from https://ddbj.nig.ac.jp/public/ddbj_database/gea/experiment/E-GEAD-000/E-GEAD-420/ on 21st May 2024. |
| Quach et al[11] | https://doi.org/10.1016/j.cell.2016.09.024 | QTD000409.all.tsv.gz, QTD000414.all.tsv.gz, QTD000419.all.tsv.gz, QTD000424.all.tsv.gz, and QTD000429.all.tsv.gz downloaded from https://www.ebi.ac.uk/eqtl/ on 13th March 2024. |
| Rizig et al[12] | https://doi.org/10.1101/2023.05.05.23289529 | Summary statistics provided by authors. |
| Shigemizu et al[13] | https://doi.org/10.1038/s41398-021-01272-3 | Summary statistics provided by authors. |

**Supplementary Table 9: List of datasets utilised in this study.**

### Supplementary Figures

#### Supplementary Figure 1

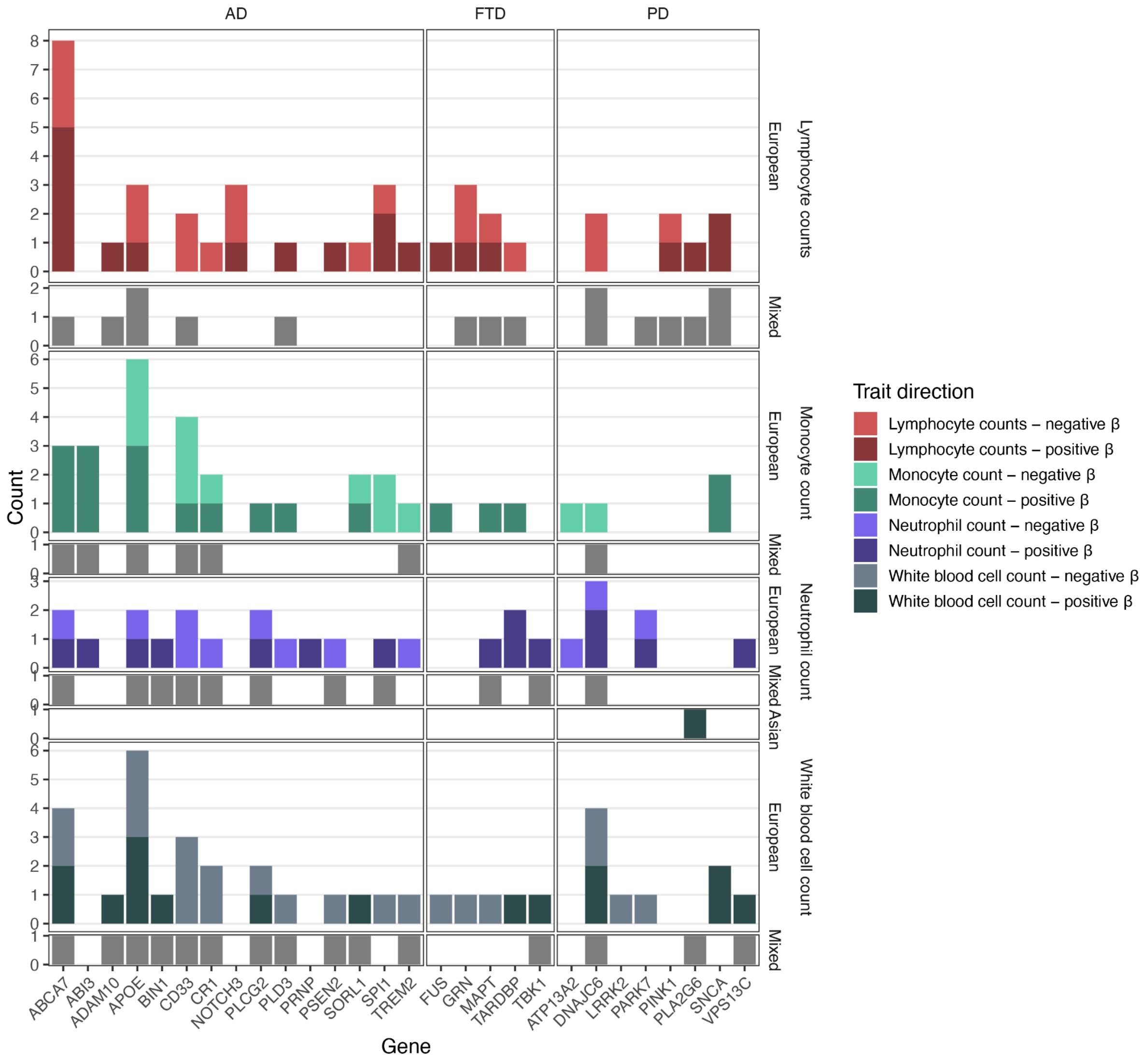


**Supplementary Fig. 1: Phenotype QTL associations between NDD genes and white blood cell count metrics.** Number of pQTL associations between NDD genes and 4 white blood cell count metrics from Chen et al[5] (top to bottom): lymphocyte, monocyte, neutrophil and total white blood cell counts. Barchart shows the count of ancestry-specific results and muti-ancestry meta-analysis. Ancestry-specific results were coloured by direction of effect (β), though in the multi-ancestry meta-analysis this was not reported.

#### Supplementary Figure 2

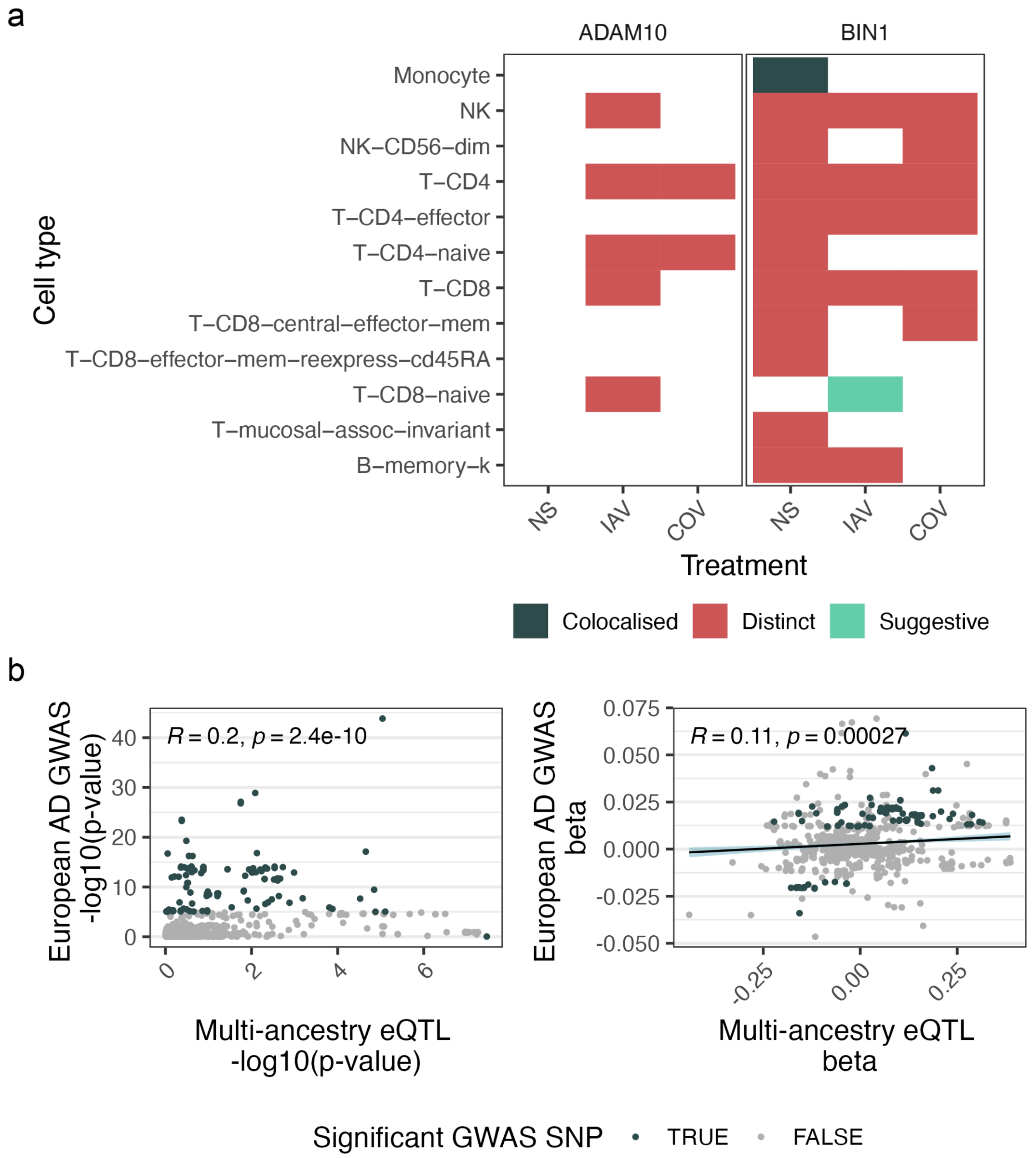


**Supplementary Fig 2. Colocalisation results of AD-causing genes in European AD GWAS and Multi-ancestry eQTL. a**, Results for genes causally implicated in AD across cell types and activation states. Colocalisation defined by PPH4 > 0.75; Distinct is defined by PPH3 > 0.75; Suggestive is defined where the ratio of PPH4/PPH3 is high (ratio_PPH4_PPH3 > $\log_2(9)$), but there is insufficient power to conclude a significant colocalisation (PPH3 + PPH4 is between 0.5-0.75). NS, no stimulation; IAV, influenza A virus; COV, SARS-CoV-2 virus). **b**, Locus analysis of significant colocalising signal at BIN1 in unstimulated monocytes,, showing p-values at left, and betas at right. Significant GWAS SNP defined by -log10(p-value) > $5x10^{-8}$, inset text shows Pearson's correlation coefficient and p-value, and beta plot also show regression line of best fit and 95% confidence intervals.

**Supplementary Figure 3**

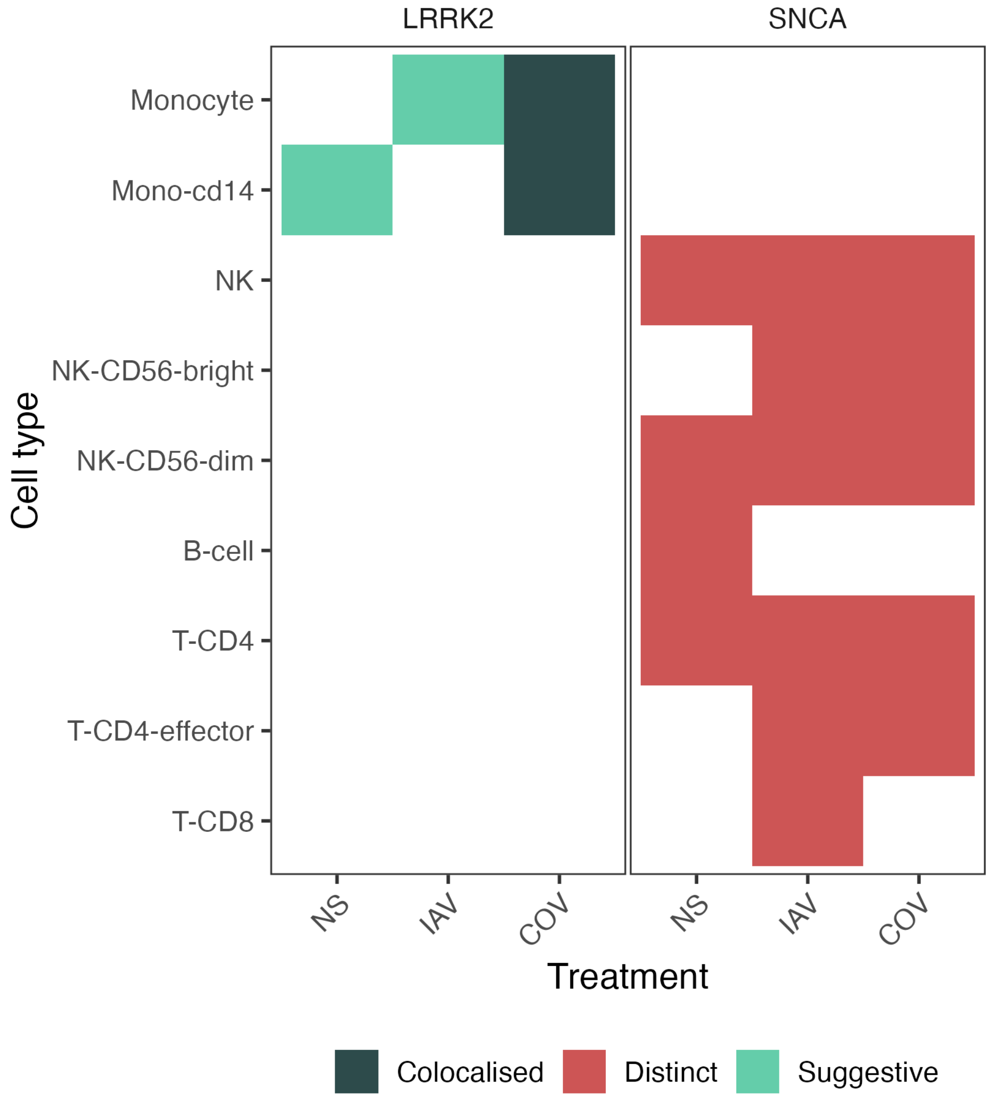


**Supplementary Fig 3. Colocalisation results in European PD GWAS and Multi-ancestry eQTL. a**, Results for genes causally implicated in PD across cell types and activation states. Colocalisation defined by PPH4 > 0.75; Distinct is defined by PPH3 > 0.75; Suggestive is defined where the ratio of PPH4/PPH3 is high (ratio_PPH4_PPH3 > $\log_2(9)$) , but there is insufficient power to conclude a significant colocalisation (PPH3 + PPH4 is between 0.5-0.75). NS, no stimulation; IAV, influenza A virus; COV, SARS-CoV-2 virus).

**Supplementary Figure 4**

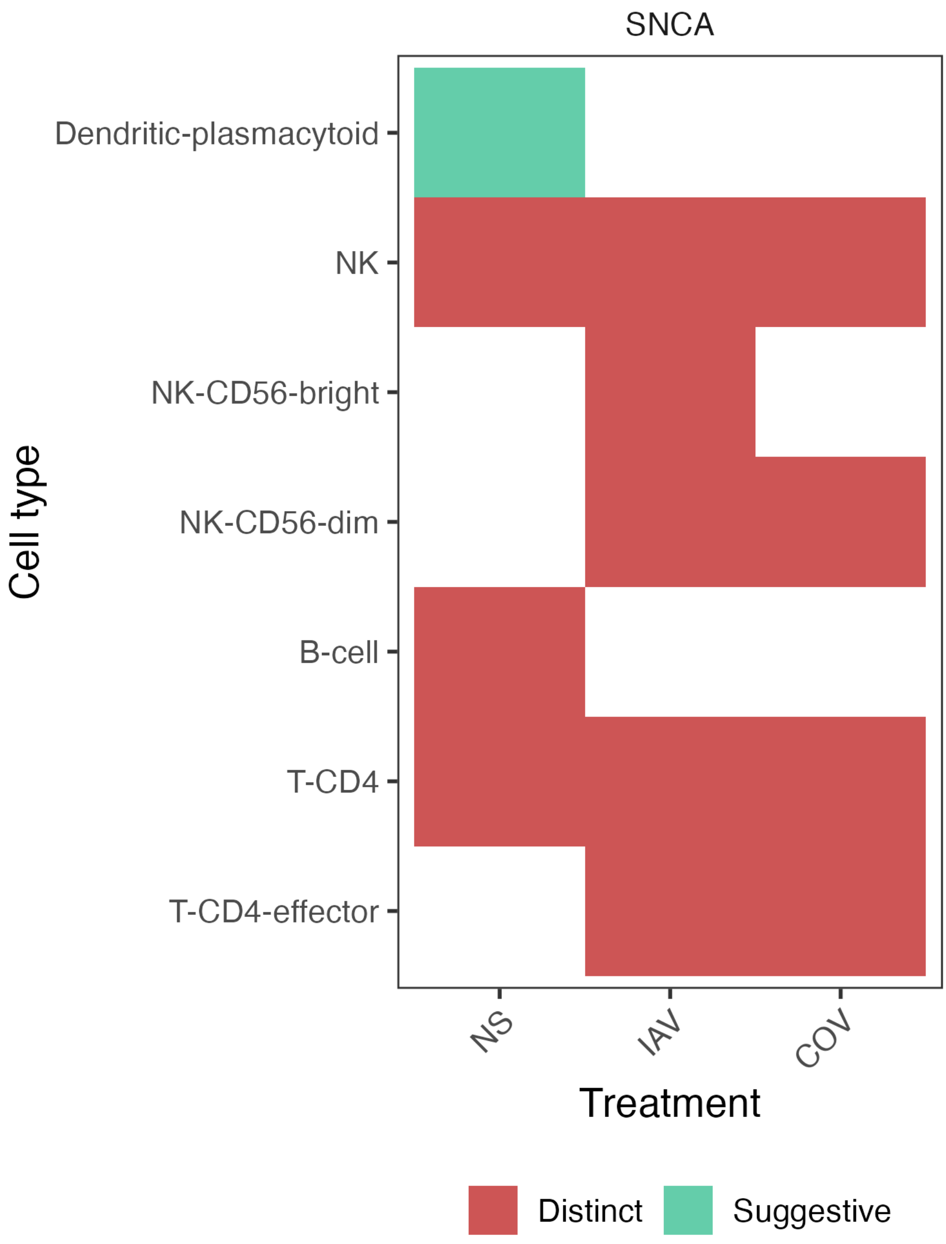


**Supplementary Fig 4. Colocalisation results in East Asian PD GWAS and Multi-ancestry eQTL. a**, Results for genes causally implicated in PD across cell types and activation states. Colocalisation defined by PPH4 > 0.75; Distinct is defined by PPH3 > 0.75; Suggestive is defined where the ratio of PPH4/PPH3 is high (ratio_PPH4_PPH3 > $\log_2(9)$) , but there is insufficient power to conclude a significant colocalisation (PPH3 + PPH4 is between 0.5-0.75). NS, no stimulation; IAV, influenza A virus; COV, SARS-CoV-2 virus.

#### Supplementary Figure 5

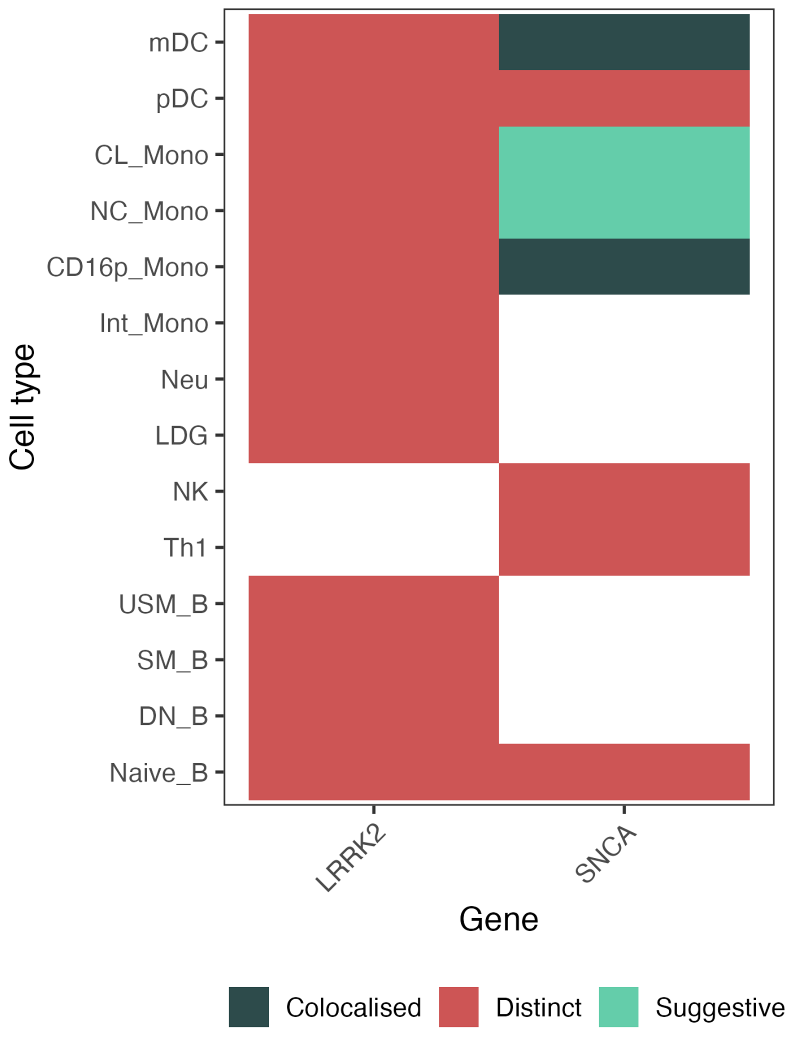


**Supplementary Fig 5. Colocalisation results in East Asian PD GWAS and East Asian eQTL.** Results for genes causally implicated in PD across cell types. Colocalisation defined by PPH4 > 0.75; Distinct is defined by PPH3 > 0.75; Suggestive is defined where the ratio of PPH4/PPH3 is high (ratio_PPH4_PPH3 > $\log_2(9)$), but there is insufficient power to conclude a significant colocalisation (PPH3 + PPH4 is between 0.5-0.75). NS, no stimulation; IAV, influenza A virus; COV, SARS-CoV-2 virus).
